# Distinct Ire1-driven transcriptional responses control morphogenesis in *Candida albicans*

**DOI:** 10.1101/2025.11.13.688365

**Authors:** Samuel Stack-Couture, Gabriela Nunes Marsiglio Librais, Bryan Lung, Julie Genereaux, Viola Halder, Vanessa Dumeaux, Rebecca S. Shapiro, Patrick Lajoie

## Abstract

The pathogenic yeast *Candida albicans* relies on morphogenesis—the transition from spherical yeast to filamentous hyphal forms—for infection. While morphogenesis requires Ire1, a transmembrane protein that canonically initiates the Unfolded Protein Response (UPR) through *HAC1* mRNA splicing, the specific mechanisms linking Ire1 to filamentation remain unclear. Using transcriptome analysis, we found that the Ire1-dependent transcriptional response driving morphogenesis is fundamentally distinct from the canonical UPR response to proteotoxic stress, with minimal overlap between programs. Remarkably, morphogenesis occurs without detectable *HAC1* splicing, and *HAC1* deletion only partially impairs filamentation, unlike complete loss with *IRE1* deletion. These findings establish that Ire1 regulates hyphal development through previously uncharacterized *HAC1*-independent pathways. Our data reveal decreased transcription of secretory proteins in an Ire1-dependent manner, providing compelling evidence that *C. albicans* possesses regulated Ire1-dependent decay (RIDD) activity—a post-transcriptional mechanism not previously characterized in this pathogen. Additionally, we identify cell wall integrity as a key *HAC1*-independent mechanism, with Ire1—but not Hac1—essential for cell wall stress tolerance and upregulation of cell wall biosynthesis genes during filamentation. Given Ire1’s essential role in pathogenesis and extensive development of Ire1-targeting compounds for mammalian systems, our findings position Ire1 as a highly promising druggable target for novel antifungal therapeutics and development of fungal-specific inhibitors.

## INTRODUCTION

Fungal disease is highly prevalent throughout the human population and causes an estimated 1.6 million deaths per year [1]. *Candida* species are one of the most common agents to cause fungal infections, with *Candida albicans* being the most common cause of invasive *Candida* infections, or candidiasis [2–5]. *Candida* infections can be life-threatening to immunocompromised individuals living with HIV or AIDS [6–9], cancer patients undergoing chemotherapy [10–12], and in transplant recipients [13–16].

*C. albicans* infection largely relies on the ability of this yeast to undergo morphogenic changes from the spherical yeast form to filamentous pseudohyphal and true hyphal forms [17,18]. The yeast form is important for dissemination through the bloodstream, whereas the filamentous forms enable *C. albicans* to escape phagocytic immune cells and penetrate host tissues [18,19]. Filamentation can be induced experimentally through various mechanisms including serum treatment, elevated temperature, nutrient starvation, chromatin accessibility and pH neutralization [2,20–25]. Changes in proteostasis also affect morphogenesis; Hsp90 inhibition, *HSF1* overexpression or depletion, and proteasome inhibition all regulate *C. albicans* filamentation [26–28]. Transcriptional control of morphogenesis is mediated by multiple pathways including the cAMP-PKA and MAPK pathways, and a pH-dependent pathway dependent on transcription factor Rim101 [20]. The cAMP-PKA pathway, mediated by both the ammonium permease Mep2 and the GPCR Grp1, is activated in response to nitrogen and amino acid starvation, high temperature, increased carbon dioxide [2,29]. cAMP-PKA activation leads to upregulation of Efg1, a key transcription factor involved in filamentation [30,31]. Mep2 also activates the MAPK pathway, which leads to transcriptional regulation of filamentation-specific genes by Cph1 [2]. Morphogenesis is also negatively regulated by transcription factors Nrg1 and Tup1 [30,32–34].

Morphogenesis in *C. albicans* requires the kinase inositol-requiring enzyme 1 (Ire1; [35–37], a transmembrane protein responsible for activating the unfolded protein response (UPR) in yeast [38–41]. The UPR is a highly conserved signaling pathway first characterized in the budding yeast *Saccharomyces cerevisiae*. Upon activation by unfolded proteins in the endoplasmic reticulum (a condition termed ER stress) or lipid bilayer stress ([42–45], Ire1 facilitates the splicing of an intron from *HAC1* mRNA [46,47]. This results in the synthesis of the Hac1 transcription factor [48–50], which translocates to the nucleus to upregulate hundreds of genes involved in processes such as ER-associated degradation, chaperone-mediated protein folding, and lipid biosynthesis, reflecting the breadth of the cellular functions regulated by the UPR [51–53].

In several species of plants, animals, and fungi, Ire1 cleaves ER-localized mRNAs and targets them for degradation through a mechanism called Regulated Ire1-Dependent Decay (RIDD) [54–57]. In *Schizosaccharomyces pombe*, there is no *HAC1* orthologue or transcriptional response to ER stress and ER stress is instead alleviated by reducing the cell’s translation burden through RIDD [58,59]. Similarly, the pathogenic yeast *Candida glabrata* mediates ER stress in an Ire1-dependent Hac1-independent manner through RIDD. *C. glabrata* does possess a *HAC1* orthologue, although *HAC1* mRNA is not spliced in response to ER stress conditions and the transcriptional response to ER stress appears to depend more on the calcineurin-Crz1 pathway [60]. The *C. albicans* ER stress response is canonically mediated by the Hac1 transcriptional response. It is unclear whether *C. albicans* possesses RIDD; however, there are several known Hac1-independent functions of Ire1 in *C. albicans*. Ire1 is required for iron uptake and virulence independent of Hac1 [61], and Ire1 attenuates ER stress by activating the HOG-MAPK pathway in a manner that may not involve Hac1 [62].

Loss of Ire1 function reduces *C. albicans* filamentation and biofilm formation, leading to decreased virulence in mouse models of disseminated candidiasis [35,36,61]. Although Ire1 is known to be required for pathogenicity in *C. albicans*, the specific target genes downstream of Ire1 involved in this process remain uncharacterized. Additionally, deleting the transcription factor Hac1 in *C. albicans* was initially shown to result in a strain that is unable to undergo filamentation [63]; however, a recent study demonstrated that Hac1 is only partially required for *C. albicans* filamentation and suggested that Hac1-independent functions of Ire1 are also required for morphogenesis [64]. Here, we performed comprehensive transcriptome analysis to characterize the specific Ire1-mediated transcriptional response required for *C. albicans* morphogenesis. We also sought to define the involvement of Hac1 in *C. albicans* morphogenesis and characterize Hac1-independent functions of Ire1 in the process.

## RESULTS

### *IRE1* is required for morphogenesis in *C. albicans*

To study the unfolded protein response, we used a strain with **<u>d</u>**iminished e**<u>x</u>**pression of *IRE1* (*ire1DX*). The *ire1DX* strain contains a deletion of one *IRE1* allele with the second *IRE1* allele placed under control of the weakly expressed *PGA5* promoter [65]. A complemented *ire1DX+IRE1-WT* strain was generated as previously described to serve as a control [36]. The *ire1DX* strain is hypersensitive to tunicamycin, an N-linked glycosylation inhibitor [66] that causes ER stress (**Fig. 1A,B**). *ire1DX* is also unable to undergo *HAC1* splicing in the presence of tunicamycin, confirming that it has dysfunctional UPR activation (**Fig. 1C**). To confirm the requirement of Ire1 in *C. albicans* morphogenesis, we performed a filamentous growth assay and found that *ire1DX* was unable to undergo filamentation when grown at 37°C in 10% fetal bovine serum (FBS) (**Fig. 1D**). *ire1DX* had a 20% and 51% decrease in mean percent filaments relative to the parental DAY286 strain and the *ire1DX+IRE1-WT* control, respectively, consistent with previous findings (**Fig. 1E**; [35,36]). Next, we examined whether the kinase and nuclease domains of Ire1 are necessary for morphogenesis by complementing the *ire1DX* strain with a previously described “kinase-dead” (KD) or “nuclease-dead” (ND) copy of *IRE1* [36]. We found that both the kinase and nuclease domain of Ire1 are required for morphogenesis, demonstrated by a 23% and a 27% decrease in mean percent filaments relative to the control, respectively (**Fig. 1F,G**). Both functions of Ire1 are also required for *C. albicans* growth on tunicamycin (**Supp Fig. 2**) [36]. Notably, despite the significant evolutionary divergence between these species, *C. albicans IRE1* was able to functionally replace *S. cerevisiae IRE1* (**Supp Fig. 3A,B**), indicating that the *HAC1* splicing capability has been evolutionarily maintained.

**Figure 1.**
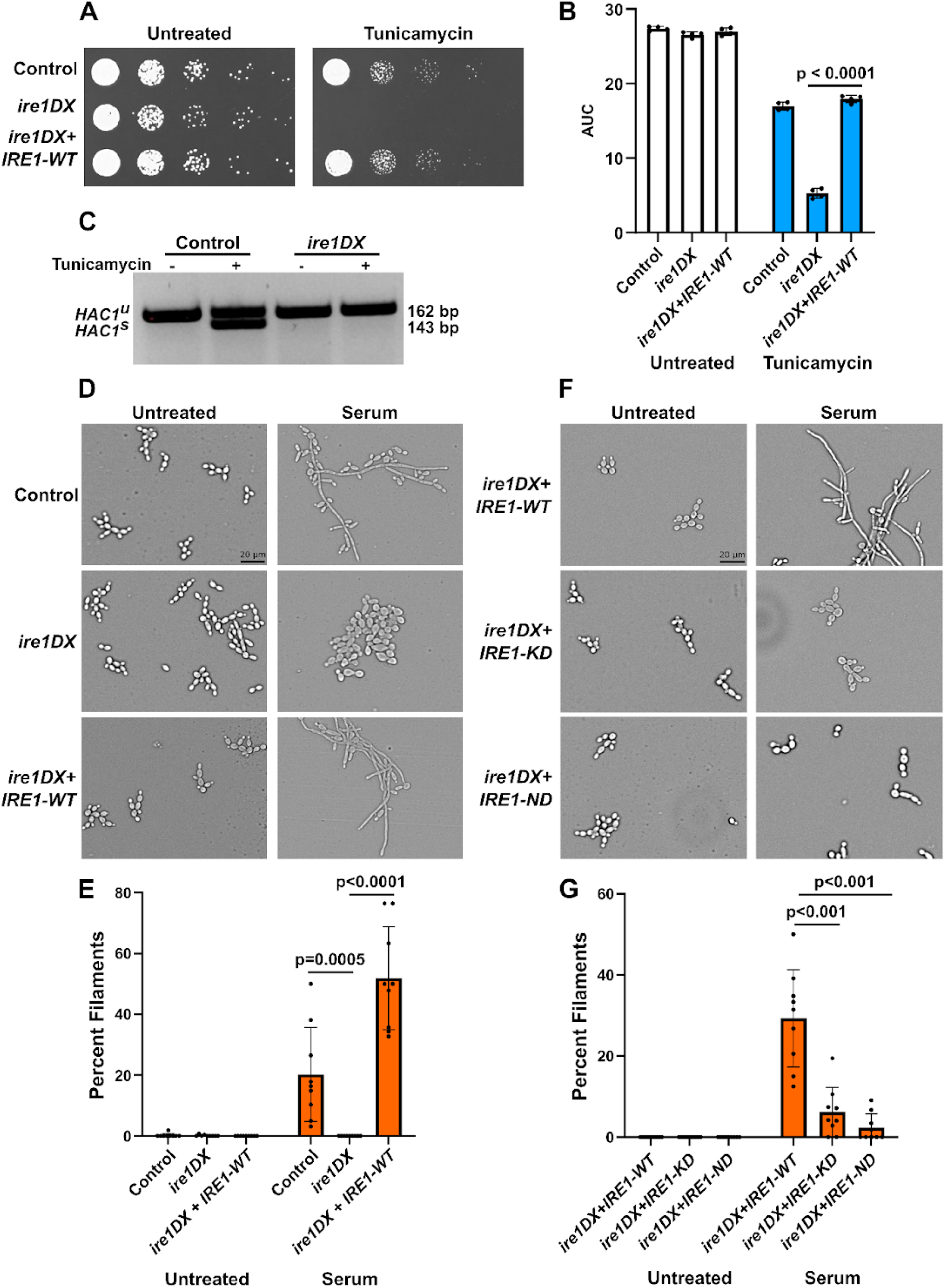
*IRE1* is required for the *C. albicans* response to the ER stressor tunicamycin and for morphogenesis. **A.** Control (DAY286), *ire1DX,* and *ire1DX+IRE1-WT C. albicans* cells were diluted to OD_600_ 0.1, serially diluted, and grown on agarose plates with the canonical ER stressor tunicamycin (1.5 µg/mL). Images acquired after 24 hours. **B.** Cells were treated with 2 µg/mL tunicamycin and grown in liquid for 24 hours. Area under the curve (AUC) was calculated and statistical analysis performed using a One-way ANOVA with Tukey’s multiple comparison test (n=4). **C.** Cells were treated with 2.5 µg/mL tunicamycin for two hours and total RNA was isolated and purified. RT-PCR was performed using primers flanking the *HAC1* intron and DNA was run on a 4% agarose gel. **D/F.** Indicated *C. albicans* strains were grown in filamentation-inducing 10% fetal bovine serum for four hours at 37°C and untreated at 30°C as a control. Images obtained using the Cytation5 cell imaging multi-mode reader. Scale bar = 20 µm, applies to all images. KD = kinase-dead; ND = nuclease-dead. **E/G.** Filamentation was quantified for 3 biological replicates. Photos were acquired from randomly selected portions of each slide, and round and filamentous cells were classified and counted by hand for each photo. Percent filaments was calculated and mean±SD was plotted.

### A distinct *IRE1* transcriptional response drives morphogenesis

Given the requirement for Ire1 during morphogenesis, we sought to determine the Ire1-regulated target genes involved in morphogenesis. Wild-type cells and *ire1DX* were treated with serum and grown at 37°C for four hours and processed for transcriptome analysis using RNA-seq. To compare the transcriptional response associated with filamentous growth to a canonical Ire1-mediated proteotoxic stress response, wild-type and *ire1DX* cells were treated with tunicamycin for two hours and similarly processed. Dimension reduction by principal component analysis showed that biological replicates for the same condition clustered close together and highlighted a treatment effect that is dependent on strain (**Fig. 2A,B**).

**Figure 2.**
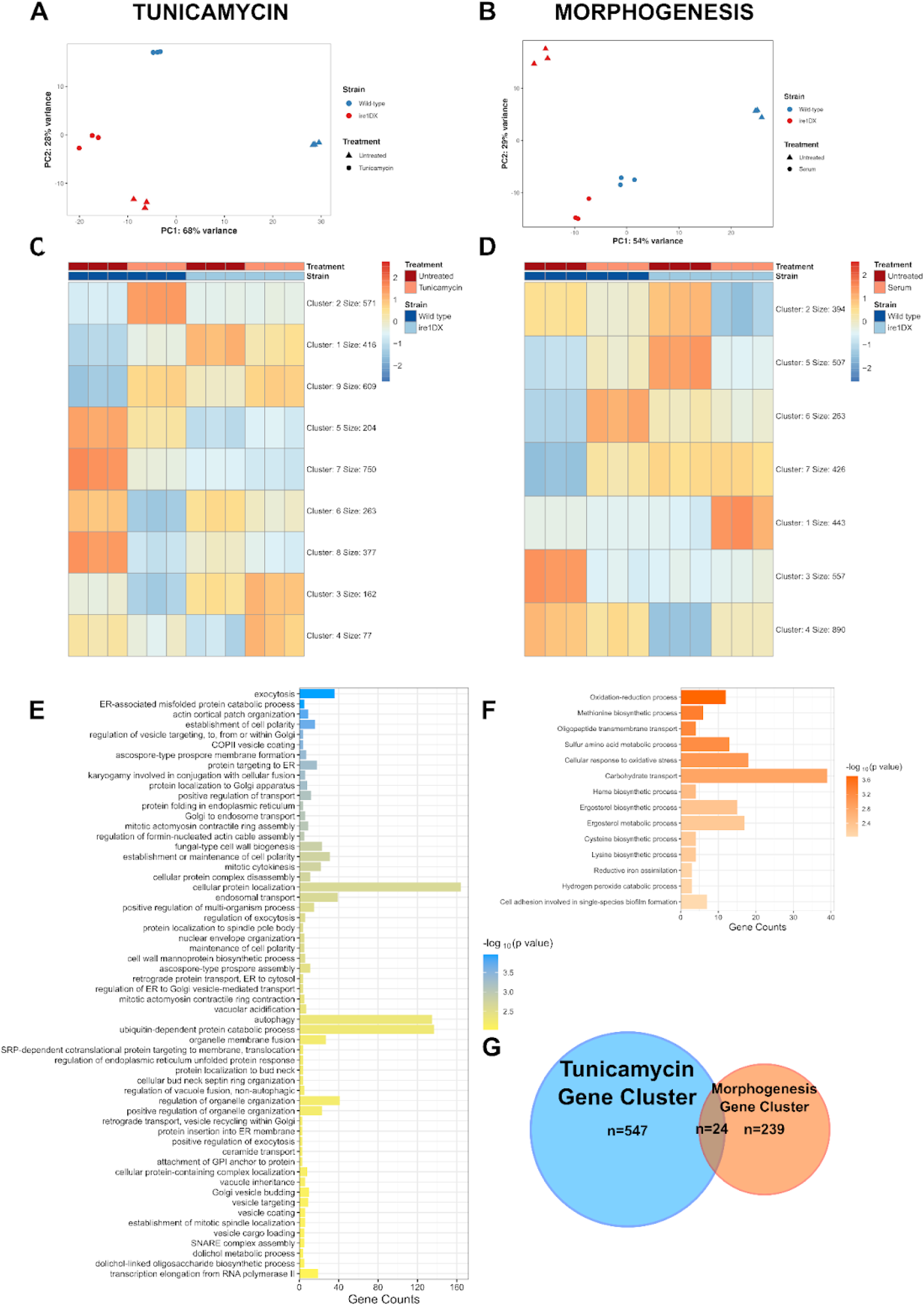
Clustering of differentially expressed genes reveals genes upregulated by Ire1 in response to tunicamycin and serum+37°C treatment. Principal component analysis of control (DAY286) and *ire1DX* treated with **A.** 1.5 μg/mL tunicamycin or **B.** 10% fetal bovine serum+37°C. Each point represents a single biological replicate (n=3). RNA sequencing data from control (DAY286) and *ire1DX* treated with **C.** tunicamycin or **D.** Serum+37°C were analyzed. Differentially expressed genes with a significant treatment*strain interaction (p_adj_<0.05) were clustered into groups based on similar expression profiles (n=3429 genes for tunicamycin and n=3480 genes for morphogenesis). Gene ontology analysis for biological processes was performed using ViSEAGO in R Studio [124] for **E.** tunicamycin cluster 2 and **F.** morphogenesis cluster 6. Only terms with p<0.01 are displayed. **G.** Venn diagram representing total number of genes in tunicamycin cluster 2 and morphogenesis cluster 6 shows little overlap.

To identify specific treatment effects dependent on strain, we identified differentially expressed genes with tunicamycin or serum+37°C treatment as a function of strain (p_adj_<0.05). Of the original 6263 genes, 3429 and 3480 genes exhibited a significant treatment*strain interaction in the tunicamycin and morphogenesis dataset, respectively. Significant genes were clustered to identify groups of genes with unique expression patterns. A group of genes (**Fig. 2C; Supp File 1**; Cluster 2, 571 genes) was upregulated in tunicamycin-treated wild-type but not in tunicamycin-treated *ire1DX;* an expression pattern expected of UPR target genes. Similarly, a group of genes (**Fig. 2D; Supp File 1**; Cluster 6, 263 genes) was upregulated in serum+37°C-treated wild-type cells but not in *ire1DX*. These results indicate that these genes capture treatment effects that are dependent on Ire1.

Gene Ontology (GO) analysis revealed that the group of genes induced by tunicamycin in an Ire1-dependent manner are enriched for biological processes known to be crucial in the UPR; this included protein targeting to ER (18 genes; p=0.00052), protein localization to Golgi apparatus (20 genes; p=0.00056), and ER-associated misfolded protein catabolic process (7 genes; p=0.00011) (**Fig. 2E**) [52,63,67]. GO analysis for the cluster of genes upregulated in an Ire1- and serum+37°C-dependent manner revealed upregulation of several genes known to form the core filamentation response in *C. albicans* (**Fig. 2F**). This includes *ALS3*, which encodes a cell wall-associated protein [68,69], and *DCK1*, which encodes a Rac1 guanine nucleotide exchange factor [70]. Interestingly, a comparison of the two Ire1-upregulated gene clusters showed that the tunicamycin and serum+37°C treatments each induced a unique transcriptional response. Only 24 genes were identified as being upregulated in both groups of genes (**Fig. 2G**). Several of these 24 genes have roles in cellular protein localization, protein folding, and post-translational modifications, as well as roles in cell wall biogenesis and amino acid metabolism (**Supp File 1**). Many of the genes localize to portions of the secretory pathway, including the ER, Golgi, and cell membrane, as well as the cell wall. This is logical given the previously highlighted role of Ire1 in cell wall homeostasis [35].

To further characterize our findings obtained through RNA sequencing, we performed RT-qPCR on several of the most differentially expressed genes from either dataset. Canonical UPR target genes *KAR2* [71] and *PDI1 [52]* were selected from the tunicamycin-upregulated genes and *ALS4* and *AHP1* were selected from the serum+37°C-upregulated genes. As expected, the expression of *KAR2* and *PDI1* was increased upon treatment with tunicamycin in wild-type but not in *ire1DX* (**Supp Fig. 4A,B**). Similarly, the expression of morphogenesis genes *AHP1* and *ALS4* increased in wild-type treated with serum+37°C but not in *ire1DX* (**Supp Fig. 4C,D**). Neither *KAR2* nor *PDI1* were upregulated in response to treatment with serum+37°C, and *AHP1* was not upregulated in response to tunicamycin, supporting the idea that the Ire1-mediated transcriptional response to both treatments varies greatly.

To determine the requirement for the identified Ire1-regulated genes in tunicamycin tolerance, 20 of the genes most differentially expressed with tunicamycin treatment in wild-type but not in *ire1DX* (Cluster 2 genes) were selected to analyze further (**Fig. 3A**). *HAC1* was not one of the top 20 hits, but given its prominent role in the UPR, it was included in the set of assessed genes [46]. Based on previous microarray data [53], 13 of these 21 genes are known UPR target genes in *S. cerevisiae* (**Supp File 1**). To obtain *C. albicans* strains with decreased expression of each of these genes, strains were selected from the publicly available *C. albicans* GRACE library (Gene Replacement and Conditional Expression) and its associated expansion library (GELC; GRACE Expanded Library Collection). The GRACE method of gene repression allows for the transcriptional repression of tetracycline promoter-regulated genes. In total, the library contains ∼3,000 mutants, with the GELC expansion adding 866 more [72,73]. Our mini-library contained four replicates of each selected strain from the GRACE and GELC libraries. The strains were grown in the presence of low (0.05 μg/mL) or high (20 μg/mL) concentrations of doxycycline to induce repression of both essential and non-essential genes, respectively, followed by a treatment with tunicamycin. Hits were identified by identifying strains with a growth defect when treated with doxycycline and tunicamycin but with no growth defect when treated with doxycycline alone (**Fig. 3B**) [74]. This screen identified several genes as being required for tunicamycin tolerance, including *HAC1*, *LHS1*, *PMI1*, *CDC11*, and *CDC12. HAC1* is a known UPR target gene required for *S. cerevisiae* to grow in the presence of tunicamycin [52,75]. *LHS1* and *PMI1* are also known UPR target genes [52,76]. *LHS1* plays a role in protein translocation into the ER and repair of misfolded proteins ([77], and *PMI1* is involved in protein glycosylation [78]. *CDC11* and *CDC12* encode septins with essential functions in cytokinesis. They have not been previously identified as UPR target genes [52]; however, ER stress is known to activate a surveillance pathway in *S. cerevisiae* termed the endoplasmic reticulum stress surveillance (ERSU) pathway. ER stress can activate this pathway, leading to mislocalization of septin subunits including Cdc11 and Cdc12 and resulting in subsequent inhibition of cytokinesis [79,80].

**Figure 3.**
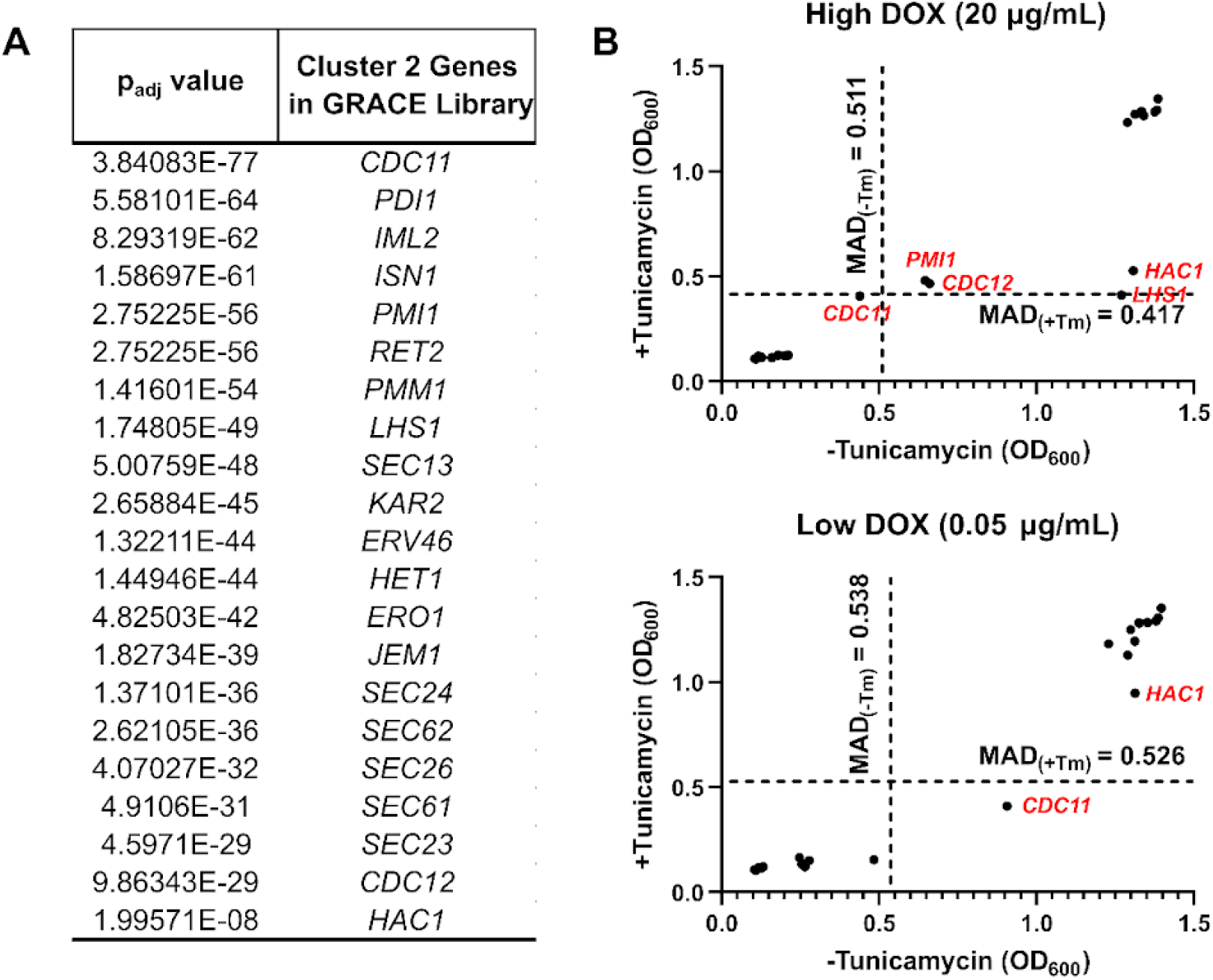
Screen of top 20 genes upregulated in an Ire1- and tunicamycin-dependent manner identifies genes required for tunicamycin tolerance. **A.** p_adj_ values for top 20 differentially expressed genes (plus *HAC1*, which was not in the top 20) in Cluster 2 of tunicamycin RNA sequencing data. Strains containing mutations of each gene were obtained from GRACE or GELC libraries [72,73]. **B.** Strains were grown overnight with low concentrations of doxycycline (DOX; 0.05 μg/mL) or high concentrations of DOX (20 μg/mL) at 30°C. Cells were then cultured in YPD containing the same concentration of DOX with or without 1.0 μg/mL tunicamycin and OD_600_ was measured after 24 hours. Data points are the average of technical quadruplicates, and dotted lines indicate the median absolute deviation (MAD) cut off values for each condition [74]. Genes of interest are highlighted in red.

We also assessed the top 20 genes upregulated in a serum+37°C- and Ire1-dependent manner (Cluster 6 genes; **Fig. 4**). A preliminary literature review was performed to determine whether these genes have known roles in morphogenesis. All but four genes had reported roles in either filamentation, biofilm formation, or both (**Supp File 1**) [33,81–89]. To further characterize these genes, PathoYeastrack was used to assess common transcription factors controlling their expression [90]. 49 transcription factors were identified as significant (p<0.00001; **Supp File 1**). The Candida Genome Database [91] was then used to characterize the regulatory functions of these transcription factors. 32 transcription factors had roles in filamentation, 24 had roles in biofilm formation, and 17 had roles in cell adhesion, whereas only 8 had roles unrelated to these functions (**Fig. 4**). Of note, Hac1 was not one of the top transcription factors identified. It is possible that this regulatory interaction has not yet been annotated in the databases referenced for the transcription factor screen; alternatively, the Ire1-dependent regulation of these genes may not involve Hac1. We decided to further examine the role of Hac1 in morphogenesis to discern this.

**Figure 4.**
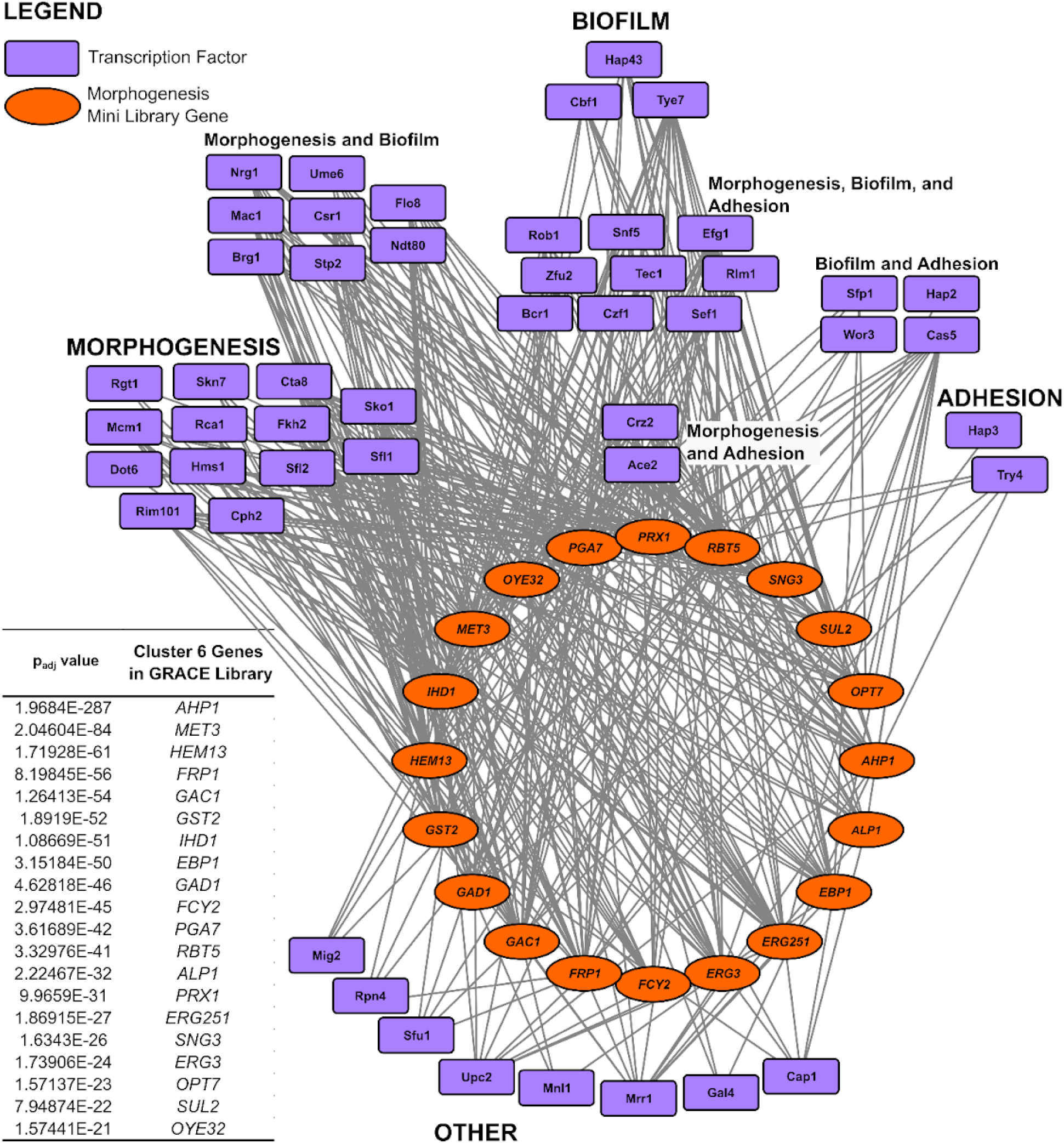
Top 20 genes upregulated in an Ire1- and serum+37°C-dependent manner (Cluster 6) are under control of morphogenesis transcription factors. The top 20 differentially expressed genes from morphogenesis Cluster 6 were subject to transcription factor analysis using PathoYeastrack. 49 transcription factors were identified; an assessment of transcription factor function using the Candida Genome Database demonstrated that most are involved in regulating morphogenesis, biofilm formation, and/or cell adhesion. Purple squares represent transcription factors; orange ellipses represent the top 20 genes. Grey edges represent interactions between transcription factors and genes. Network created in Cytoscape. p_adj_ values for top 20 differentially expressed genes in Cluster 6 of morphogenesis RNA sequencing data shown on bottom left.

### Morphogenesis is not associated with *HAC1* splicing

Given the diverging Ire1-regulated transcriptional responses to tunicamycin and filamentation-inducing conditions, we investigated the requirement of *HAC1* for morphogenesis. For this, we developed both *ire1Δ/Δ* and *hac1Δ/Δ* strains using a CRISPR-Cas9 based deletion protocol optimized for *C. albicans [92]*. DAY286 carrying Cas9 with no guide RNA (DAY286-Cas9) served as the control.

Next, we performed a morphogenesis assay and found that while *ire1Δ/Δ* lost the ability to transition to the filamentous form with serum+37°C (0.88% filaments), *hac1Δ/Δ* could still form filaments (11.30% filaments) but to a much lesser degree than the control (87.95% filaments; **Fig 5A/B)**. These *hac1Δ/Δ* filaments appeared much shorter than those of the control, and they did not clump together to form large flocs, as is typical with wild-type cells grown under filamentation-inducing conditions [93] (**Supp Fig. 5**). This suggests that while *IRE1* is required for morphogenesis, *HAC1* is only partially required. This finding is consistent with previously reported results that show that a *hac1Δ/Δ* strain grown in liquid Spider medium has a filamentation defect that is less pronounced than an *ire1Δ/Δ* strain [64].

**Figure 5.**
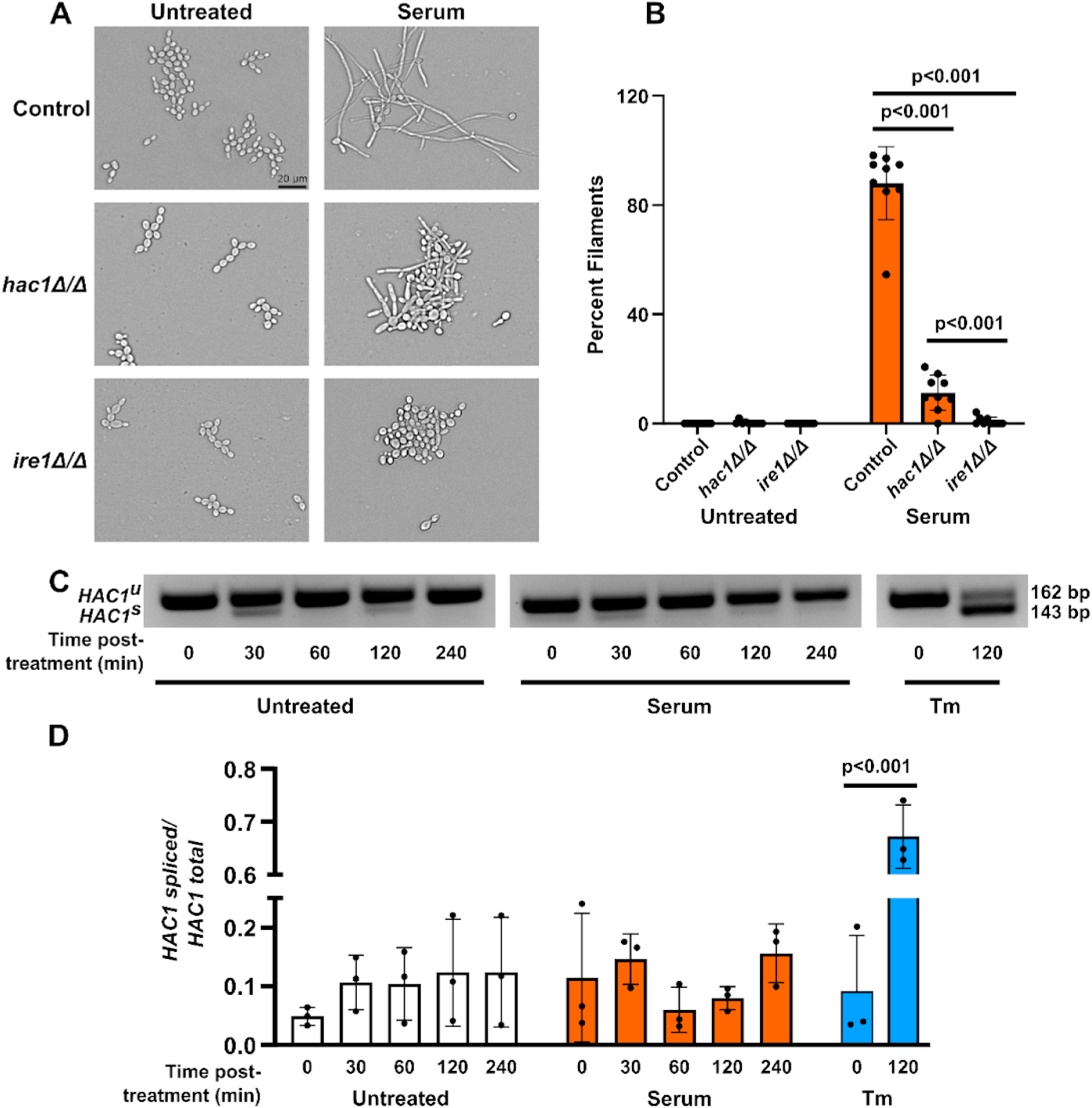
*HAC1* splicing is not required for morphogenesis. **A.** Control (DAY286-Cas9), *hac1Δ/Δ*, and *ire1Δ/Δ C. albicans* cells were grown in filamentation-inducing 10% fetal bovine serum for four hours at 37°C and untreated at 30°C as a control. Images obtained using the Cytation5 cell imaging multi-mode reader. Scale bar = 20 µm, applies to all images. **B.** Filamentation was quantified for 3 biological replicates. Photos were acquired from randomly selected portions of each slide, and round and filamentous cells were counted by hand for each photo. Percent filaments was calculated and mean±SD was plotted. **C.** Control (DAY286) cells were grown for two hours, then treated with 10% serum at 37°C or 1.5 μg/mL tunicamycin at 30°C. Total RNA was isolated from cells at the indicated time points after treatment, and RT-PCR was performed using primers flanking the *HAC1* intron. DNA was run on a 4% agarose gel. **D.** Bands were quantified using ImageJ [114] to calculate the mean grey value (MGV), and data are presented as a ratio of spliced *HAC1* to total *HAC1* (spliced + unspliced bands). Statistical analysis performed in SPSS using a One-way ANOVA with Tukey’s multiple comparison, n=3, mean±SD plotted.

To determine if *HAC1* splicing occurs in response to filamentation-inducing conditions, cells were treated with serum+37°C and *HAC1* splicing was assessed over four hours. As a control, cells treated with tunicamycin showed significant *HAC1* splicing after 120 minutes. Conversely, cells treated with serum+37°C showed no detectable *HAC1* splicing even after 240 minutes (**Fig. 5C,D**). This suggests that Ire1 is likely regulating morphogenesis through a mechanism that is independent of *HAC1* splicing.

### Ire1-dependent downregulated genes are enriched in secretory proteins

We next investigated the mechanism by which Ire1 may be regulating morphogenesis independently of Hac1. In several organisms, Ire1 functions independently of Hac1 through regulated Ire1-dependent decay (RIDD), where ER-localized mRNAs are cleaved by Ire1 and subsequently degraded [54–57]. While RIDD is a Hac1-independent process of Ire1, it is unclear whether *C. albicans* possesses RIDD. Given that morphogenesis required the Ire1 ribonuclease domain but did not induce *HAC1* splicing, we decided to probe our RNA sequencing data to look for evidence of RIDD.

To identify possible RIDD targets, we searched for transcripts that were downregulated upon treatment with tunicamycin or serum+37°C in wild-type cells but not in *ire1DX*, following a similar analysis performed in *Schizosaccharomyces pombe* [58]. To perform this analysis, we studied the effect of strain in the treated samples (Tunicamycin or Serum+37°C) and the treatment effect in the wild-type strain. Genes with a significant fold change (p_adj_<0.05) for both variables were plotted (**Fig. 6A,B**). 30 genes had a decrease in expression greater than 1.5-fold associated with both strain and tunicamycin treatment (**Fig. 6A**, blue dots; **Supp File 1**). *PDI1* and *KAR2*, known UPR target genes [52,71], were identified as having an increase in expression greater than 1.5-fold associated with both strain and treatment (**Fig. 6A**, pink dots). 23 genes had a decrease in expression greater than 1.0-fold associated with both strain and serum+37°C treatment (**Fig. 6B**, orange dots; **Supp File 1**).

**Figure 6.**
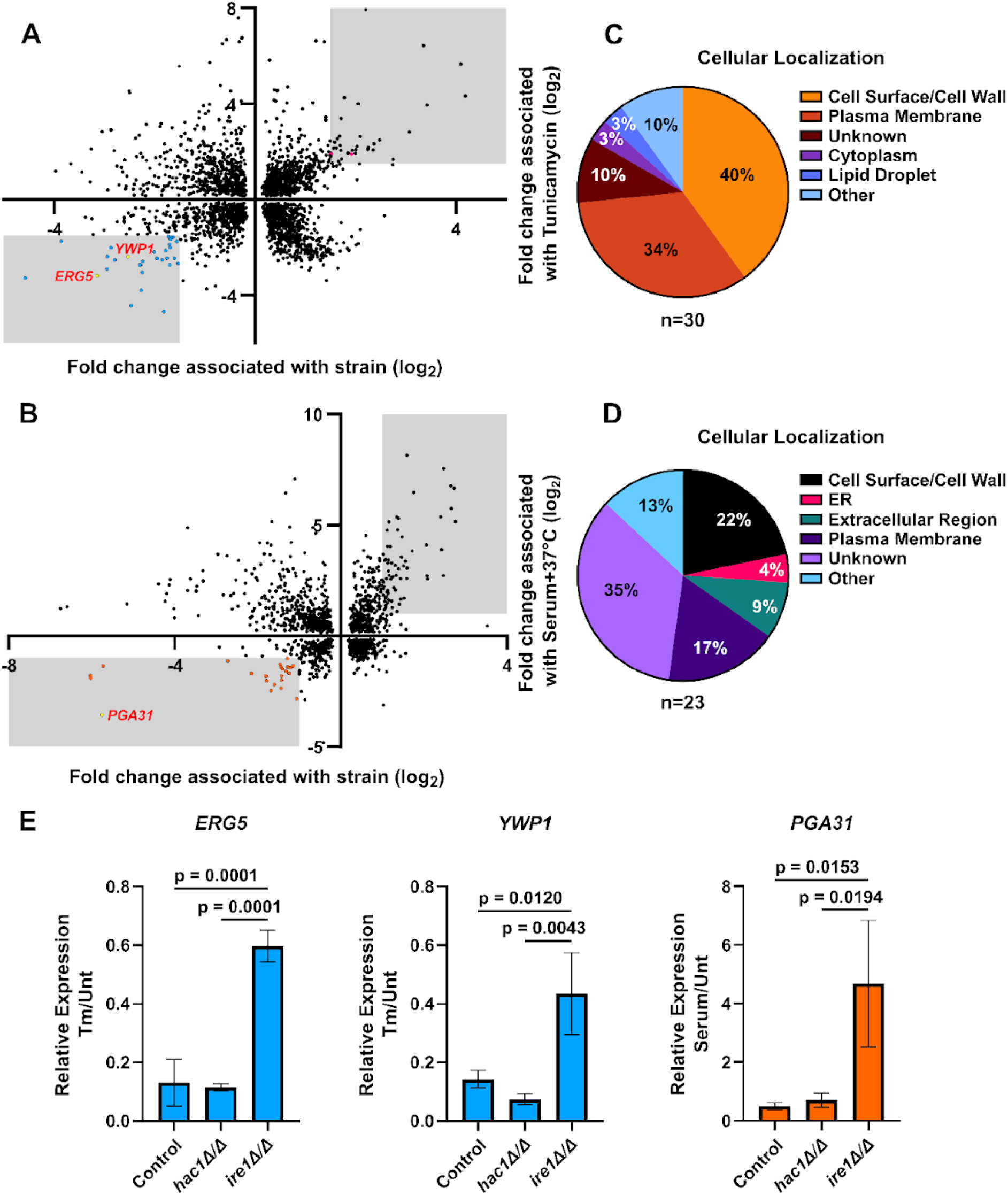
Genes with decreased expression associated with strain and tunicamycin or serum+37°C treatment encode predominantly secretory proteins. RNA sequencing data was analyzed for genes with decreased expression associated with strain and **A** tunicamycin or **B** serum+37°C treatment. All genes plotted have a significant fold change for both variables (p_adj_<0.05). Blue dots represent genes with a decrease in expression greater than 1.5-fold associated with both strain and tunicamycin treatment (n=30). Orange dots represent genes with a decrease in expression greater than 1.0-fold associated with both strain and serum+37°C treatment (n=23). Pink dots represent *PDI1* and *KAR2*, known UPR target genes. Yellow dots with labels correspond to the genes in 6E. The cellular localization of each gene’s protein product was identified using the Candida Genome Database for the **C** tunicamycin and **D** morphogenesis datasets. **E.** Relative gene expression for *ERG5* and *YWP1* in tunicamycin-treated cells and *PGA31* in serum+37°C-treated cells was assessed using RT-qPCR. Control is DAY286-Cas9. For each strain, a relative expression ratio of treated (tunicamycin or serum+37°C) to untreated cells was calculated using 2^-ΔΔCt^ values. n=3, mean± SD shown, one-way ANOVA with Tukey’s multiple comparisons performed in GraphPad Prism.

RIDD targets are primarily secretory proteins produced in the endoplasmic reticulum [55,57,58]. To characterize the transcripts with decreased expression associated with strain and tunicamycin treatment, the Candida Genome Database was used to identify the protein product and cell component of each gene [91]. Analysis of the cellular compartment of each gene’s protein product revealed an enrichment in secretory proteins; 74% of the proteins localize to the cell membrane, cell wall, or cell surface (**Fig. 6C**). Some of these proteins include GPI-linked cell wall proteins, membrane transport proteins, and several membrane-localized ferroxidases. Additionally, 36.7% of these genes were predicted to have a transmembrane domain based on sequence homology. Several genes encoding non-secretory pathway proteins were identified; these genes encode proteins localized to the nucleus, cytoplasm, lipid droplets, and one gene that encodes for a portion of the snoRNP complex (**Supp File 1**). A similar analysis was performed for genes with decreased expression associated with strain and serum+37°C treatment, and a similar trend was found. 52% of those genes encode for proteins that localize to the cell surface, cell membrane, ER, or extracellular region, and 26% were predicted to have a transmembrane domain (**Fig. 6D; Supp File 1**).

To validate potential Ire1 RIDD activity, we assessed transcript levels of candidate genes in wild-type, *hac1Δ/Δ*, and *ire1Δ/Δ* strains under both tunicamycin and filamentation-inducing conditions (**Fig. 6E**). In wild-type cells treated with tunicamycin, *ERG5* and *YWP1* transcripts showed the expected decrease in the treated versus control ratio, while *PGA31* decreased under filamentation conditions, consistent with Ire1-mediated mRNA degradation. Importantly, these decreases were absent in *ire1Δ/Δ* cells for both treatments, confirming that the transcript reduction is dependent on Ire1 function, while *hac1Δ/Δ* cells showed similar reductions to wild-type, supporting a Hac1-independent RIDD mechanism.

### A Hac1-independent function of Ire1 regulates the response to cell wall stress

We next explored another mechanism by which Ire1 may be regulating morphogenesis independently of Hac1. We assessed genes upregulated in an Ire1- and serum+37°C-dependent manner (Cluster 6 genes) using Gene Ontology analysis for cellular compartment and identified an enrichment in genes encoding cell wall proteins (**Supp File 1**).

Cell wall protein production is known to increase during morphogenesis to support growth at the hyphal tip [94,95]. Cell wall proteins are also linked to Ire1 and the UPR through the cell wall integrity pathway (CWI). The CWI is a stress pathway responsible for resolving cell wall damage [96]. During ER stress, the transport of proteins to the cell wall can become impaired and cause cell wall perturbations, leading to cell wall stress. Alternatively, cell wall stress can cause the cell to increase its protein folding capacity in an attempt to alleviate cell wall damage, leading to increased ER stress [97]. Given the known importance of Ire1 in both morphogenesis and cell wall integrity [35,36,98], we investigated whether morphogenesis could be rescued in *ire1Δ/Δ* by treatment with the osmotic stabilizer sorbitol. Sorbitol has been shown to rescue cell wall defects in various yeast [36,99,100]. We did not observe a significant increase in percent filaments when *ire1Δ/Δ* was grown with sorbitol (**Supp Fig. 6**).

Next, we sought to determine if Hac1 is required for the Ire1-dependent regulation of cell wall integrity. When grown on the cell wall stressors calcofluor white and caspofungin, *hac1Δ/Δ* grew less than the Cas9-control but more than *ire1Δ/Δ* (**Fig. 7A**). We also assessed the expression of cell wall genes in *hac1Δ/Δ* and *ire1Δ/Δ* under filamentation conditions. *PGA7*, *RBT5*, and *IHD1* were selected from the group of genes upregulated in an Ire1- and serum+37°C-dependent manner. All three encode glycophosphatidylinositol (GPI)-linked cell wall proteins and are known to be upregulated in response to filamentation cues [101]. We identified a similar trend; *IRE1* was required for the expression of all three genes under morphogenesis-inducing conditions, whereas *HAC1* was either partially required (*PGA7*, *RBT5*) or not required at all (*IHD1*; **Fig. 7B**).

**Figure 7.**
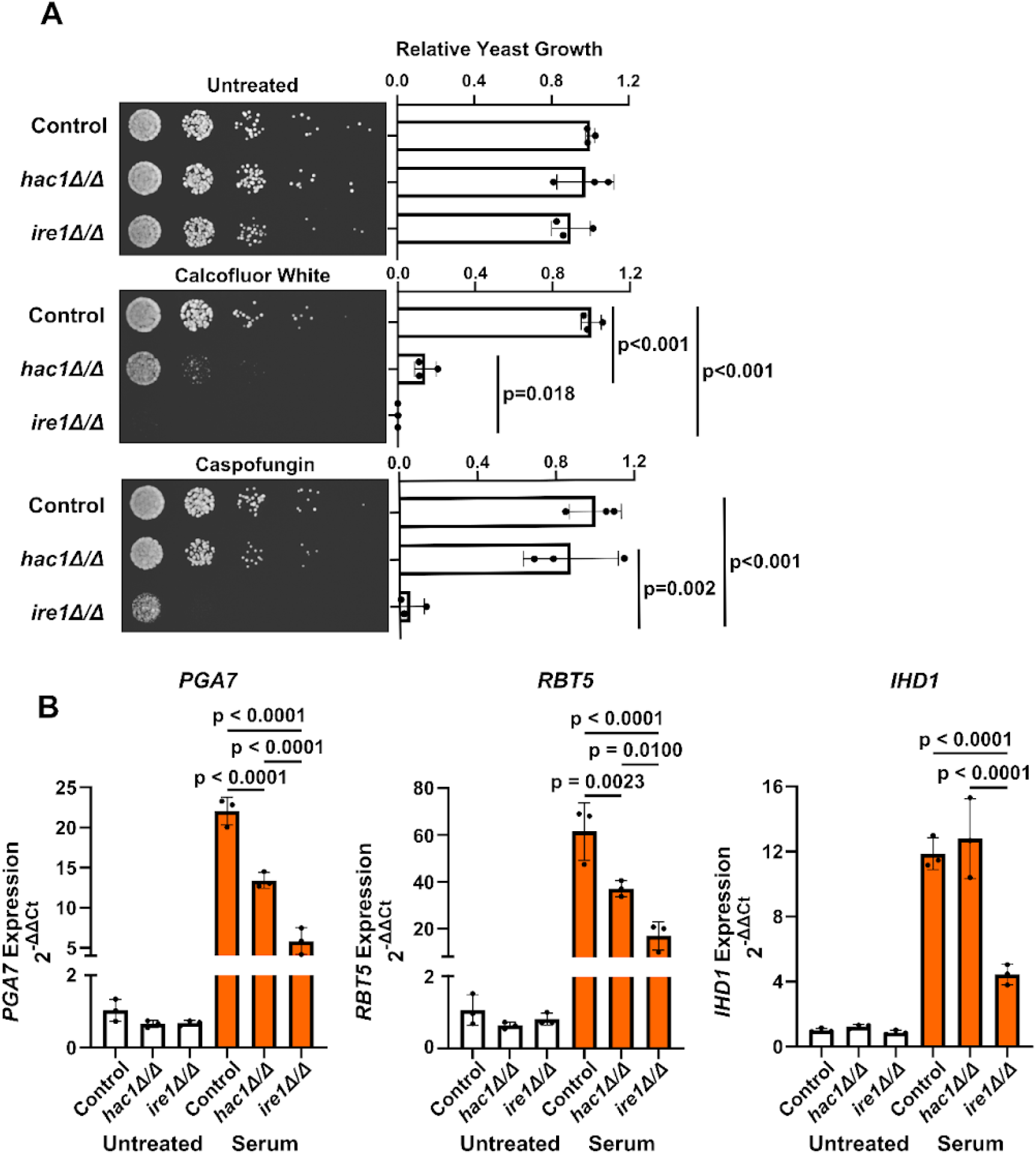
*HAC1* is not required for cell wall stress to the same degree as *IRE1*. **A.** Control (DAY286-Cas9), *hac1Δ/Δ*, and *ire1Δ/Δ C. albicans* cells were spotted on plates containing 10 µg/mL calcofluor white and 0.01 µg/mL caspofungin. Images acquired after 24 hours. Statistical analysis performed using a One-way ANOVA with Tukey’s multiple comparison, n=3, mean±SD plotted. **B.** Relative expression of GPI-anchored cell wall genes. One-way ANOVA with Tukey’s multiple comparisons. Mean±SD shown, n=3.

## DISCUSSION

### Hac1-independent regulation of hyphal growth by Ire1

While our results confirm the canonical role of Ire1 in activating the UPR through Hac1-mediated transcription during ER stress, the stark disconnect between Ire1’s essential role in morphogenesis and Hac1’s partial requirement suggests that alternative Ire1-dependent pathways drive filamentous growth in *C. albicans*. The absence of detectable *HAC1* splicing during serum+37°C-induced filamentation, coupled with the distinct transcriptional signatures during ER stress and morphogenesis conditions, underscores the critical need to identify the Hac1-independent mechanisms through which Ire1 regulates the yeast-to-hyphal transition. Previous work has demonstrated that the UPR program displays remarkable plasticity, with differential target gene expression patterns that are specifically tailored to the type of cellular stress rather than simply providing blanket upregulation of all UPR targets [53]. This stress-specific modulation of UPR target genes suggests that the network can be remodeled differentially according to the specific functional needs of the cell, which would explain why proteotoxic stress and morphogenesis-inducing conditions elicit distinct transcriptional responses despite both involving activation of the canonical UPR mediator Ire1.

Our findings demonstrating the requirement for a functional Ire1 in *C. albicans* morphogenesis are in accordance with previously reported data [35–37]. Recent studies have also demonstrated differing roles for Ire1 and Hac1 in liquid filamentation with Spider media and colony formation on solid medium. Deletion of *HAC1* decreased filamentous growth in liquid media, but not to the extent that a deletion of *IRE1* did [64]. Additionally, while *IRE1* was found to be required for inhibiting wrinkly colony formation on solid media, *HAC1* only prevented wrinkly colony formation for 48 hours. After 72 hours, colonies became wrinkly, suggesting that *HAC1* is only partially required and that alternative mechanisms downstream of Ire1 are involved in inhibiting wrinkly colony formation on solid media [64]. Several other Hac1-independent roles of Ire1 have been identified in *C. albicans*. Ire1 is required for the localization of iron permease Ftr1 to the plasma membrane, whereas Hac1 is dispensable [61].

And similarly, while Ire1 is needed for the tolerance of elevated concentrations of manganese, inactivation of *HAC1* does not influence manganese tolerance [102]. In several other fungi, Ire1 functions independently of Hac1 through RIDD [54]. *C. glabrata* Ire1 possesses kinase and ribonuclease activity similar to *C. albicans* Ire1. However, *C. glabrata* Ire1 does not facilitate *HAC1* splicing. Although this yeast has a *HAC1* orthologue, the transcriptional response to ER stress relies on calcium signaling and the Slt2 MAPK pathway [54,60]. Ire1 instead facilitates RIDD independently of *HAC1* [60]. Our finding that morphogenesis requires the Ire1 ribonuclease domain but does not induce *HAC1* splicing led us to question whether *C. albicans* possesses RIDD (**Fig. 1F and 5C,D**).

In other fungi such as the fission yeast *Schizosaccharomyces pombe*, which lacks *HAC1* and relies solely on RIDD for ER stress alleviation, RIDD targets primarily encode for proteins targeted to the ER [54,55,58]. Similarly, in *C. glabrata*, the majority of targets encoded GPI-anchored cell wall and membrane proteins [60]. In accordance with this, over 70% of the genes we identified as having decreased expression associated with strain and tunicamycin treatment encode for proteins localizing to the cell surface/cell wall and the plasma membrane (**Fig. 6A,C**), and a similar pattern exists for serum+37°C treatment (**Fig. 6B,D**). Thus, our data suggest *C. albicans* may possess a previously unidentified RIDD mechanism. The primary function of RIDD is to decrease the mRNA load in the ER, thereby reducing the overall translational burden in the cell and enabling a quick return to ER homeostasis [40,103]. RIDD is therefore beneficial in the response to ER stressors like tunicamycin which increases protein misfolding by disrupting N-linked glycosylation [104,105]. However, the role of RIDD in responding to serum+37°C-induced morphogenesis is less clear. We observed no *HAC1* splicing during filamentation, suggesting that cells are not experiencing ER stress and therefore RIDD is not functioning to relieve ER stress through decreased translation of secretory proteins. It was previously demonstrated that many genes involved in pathogenesis and filamentation show reduced translational efficiency despite transcriptional induction during *C. albicans* morphogenesis, suggesting widespread translational fine-tuning mechanisms are at play [94]. Based on these findings, RIDD could facilitate morphogenesis by selectively degrading specific secretory pathway mRNAs to redirect cellular resources toward the energy-intensive processes required for hyphal growth, such as rapid cell wall expansion and polarized development. This mechanism would also help manage ER stress during the extensive cellular remodeling of the yeast-to-hyphal transition, while providing precise post-transcriptional control over protein expression timing and magnitude to complement the transcriptional programs driving morphogenesis.

Finally, RIDD targets are frequently found to share similar sequences. Mammalian RIDD targets possess a CUGCAG consensus sequence cleaved by Ire1 that’s accompanied by a stem-loop structure that is also found in *XBP1* [40,106,107]. In *S. pombe*, RIDD targets share a UGC consensus sequence where Ire1 cleavage of the transcript occurs [58]. However, in *C. glabrata*, the question of a consensus sequence amongst RIDD targets remains to be determined [60]. Future studies should examine whether the targets identified here and in other fungi possess a consensus sequence.

### Cell wall integrity and morphogenesis

In addition to RIDD, we also explored cell wall stress as a possible connection between Ire1 and morphogenesis given that Ire1 was previously found to be included in the circuitry that controls cell wall homeostasis in *C. albicans* [35]. Our RNA sequencing revealed an abundance of cell wall genes upregulated during morphogenesis, and RT-qPCR confirmed a requirement for *IRE1* but not *HAC1* in the upregulation of these genes. Similarly, *IRE1* but not *HAC1* was required for cell growth in the presence of cell wall stressors. A previous study supports these findings; a strain with decreased expression of *IRE1* was sensitive to cell wall stressor calcofluor white but did not undergo *HAC1* splicing [36]. Cell wall integrity relies on Ire1 and the UPR because the ER is the primary site of cell wall protein production. Impaired Ire1 function thus disrupts cell wall protein synthesis and can lead to decreased cell wall integrity and stress [35,97,98]. This may explain why we observe impaired filamentation in *IRE1*-deficient strains; the morphological transition from yeast to hyphae relies on the rapid expansion of the cell wall at the hyphal tip, something that a strain lacking Ire1 may be unequipped to handle [95]. However, it is important to note that deleting *HAC1* did not have as detrimental an effect on filamentation as deleting *IRE1* did, suggesting that the UPR target gene upregulation is not the primary Ire1-dependent mechanism responsible for morphogenesis. Perhaps Ire1 interacts with other downstream targets to facilitate adaptation to cell wall stress. It would be of future interest to determine if there are any direct protein-protein interactions between Ire1 and cell wall stress-related signaling proteins.

Our findings align with recent work demonstrating that Ire1 maintains basal ER functionality independent of Hac1 activation [61]. It was shown that Ire1 is essential for proper localization of the iron permease Ftr1 to the plasma membrane, and that *ire1Δ* mutants exhibit Ftr1 retention in the ER despite the absence of detectable ER stress or *HAC1* splicing during iron limitation. This quality control function of Ire1 may be particularly critical during morphogenesis, when the extensive cellular remodeling and rapid cell wall expansion at the hyphal tip require efficient trafficking of secretory pathway proteins—including the cell wall proteins we identified as Ire1-dependent targets. The inability of *ire1*Δ but not *hac1*Δ mutants to properly upregulate cell wall biosynthesis genes during filamentation suggests that Ire1’s role in maintaining secretory pathway homeostasis is essential for supporting the massive protein flux required for hyphal development, even in the absence of overt ER stress (**Fig. 8**).

**Figure 8:**
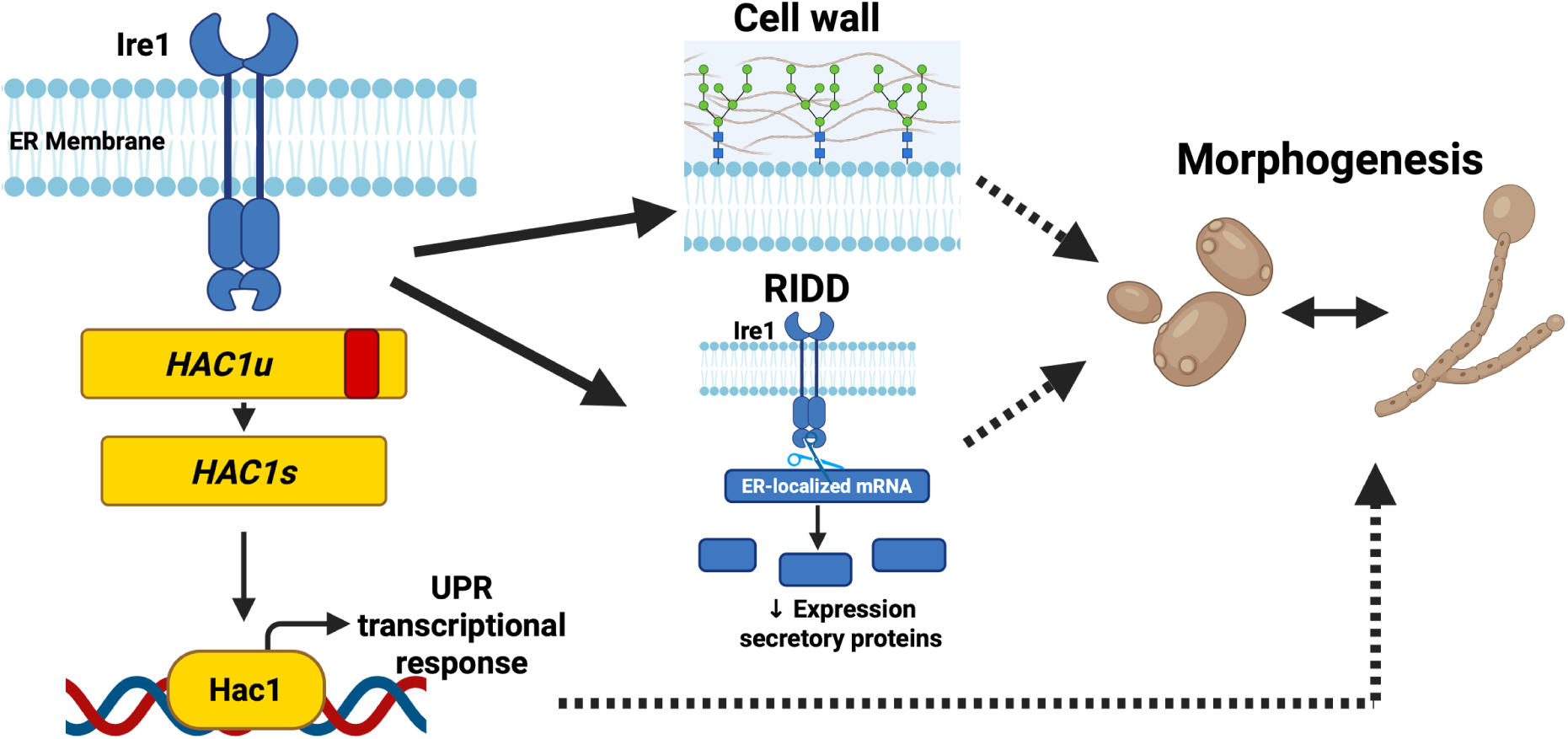
A distinct Ire1-driven transcriptional response controls morphogenesis in *C. albicans*. Model depicting the multiple mechanisms by which Ire1 regulates morphogenesis in *C. albicans*. Unlike the canonical ER stress response, filamentation-inducing conditions do not trigger *HAC1* splicing, yet require Ire1 for hyphal development. Ire1 regulates morphogenesis through three potential mechanisms: (1) Hac1-independent regulation of cell wall integrity genes, including upregulation of cell wall proteins and cell wall stress tolerance; (2) Regulated Ire1-dependent decay (RIDD) of mRNAs encoding secretory pathway proteins, particularly those localizing to the cell membrane and cell wall; and (3) a Hac1-dependent response that is only partially required for filamentation. The kinase and nuclease domains of Ire1 are both essential for morphogenesis. This model highlights the mechanistic divergence between the Ire1-dependent transcriptional programs activated during proteotoxic stress versus those required for morphogenesis, with minimal overlap between the two responses. Created in BioRender. Lajoie, P. (2025) https://BioRender.com/f97luas.

### Perspectives and Considerations

It would be of interest to explore whether the requirement for Ire1 but not Hac1 in morphogenesis persists with other filamentation-inducing conditions. Our research utilized fetal bovine serum and an elevated temperature of 37°C to induce filamentation. Serum is thought to induce filamentation through the breakdown of serum glycoproteins which yields N-acetylglucosamine (GlcNAc) and proline, both of which can independently induce filamentation [20]. Elevated temperature removes Hsp90 inhibition of Ras1, leading to activation of the cAMP-PKA and subsequent filamentation mediated by the transcription factor Efg1 [20,26]. Filamentation can also be induced through other means, including amino acid starvation, pH changes, elevated carbon dioxide, and internalization by host immune cells [2,20]. It has previously been established that the transcriptional responses to different filamentation-inducing conditions vary greatly [101]. Additionally, a recent study demonstrates differential requirements for the cAMP-PKA pathway *in vivo* compared to *in vitro*; deleting adenylyl cyclase *CYR1* and either catalytic subunit of PKA (*TPK1* and *TPK2*) resulted in minimal changes to filamentation of cells injected into the ears of live mice, whereas the same deletions caused significant decreases in filamentation for cells grown *in vitro* [108]. The requirement for certain genes for filamentation *in vivo* and *in vitro* can also vary in a strain-dependent manner [109,110]. Given the variation that can be observed in *C. albicans* filamentation, it’s possible that this Hac1-independent role of Ire1 in regulating morphogenesis is not observable when cells are grown in different filamentation-inducing conditions. Future studies should investigate this possibility.

Given that Ire1 is essential for *C. albicans* morphogenesis and pathogenesis, and considering the extensive development of *IRE1*-targeting small molecules for mammalian systems—including both RNase inhibitors and kinase modulators— [111] Ire1 emerges as a highly druggable target for antifungal therapy. The opportunity exists both to repurpose existing mammalian UPR modulators and to leverage AI-driven drug discovery approaches to develop yeast-specific Ire1 inhibitors that could serve as promising combination therapies with current antifungal agents, potentially overcoming resistance mechanisms and enhancing therapeutic efficacy against invasive candidiasis and other fungal diseases [112,113].

## MATERIAL AND METHODS

### Yeast Strains and Cell Cultures

All *C. albicans* strains used in this study are listed in **Supp File 1**. All strains were grown from frozen glycerol stocks on YPD (1% yeast extract, 2% peptone, 2% dextrose) or selective synthetic complete (SC; with appropriate amino acids) media plates for 48 hours at 30°C. All liquid cultures were grown in triplicate overnight in 5 mL of YPD or SC media at 30°C in a rotating drum.

### Reagents

Stock solutions of tunicamycin (5 mg/mL in DMSO; Millipore Sigma, Burlington, MA, USA), fetal bovine serum (100%; Millipore Sigma), caspofungin (1 mg/mL in H_2_O; Millipore Sigma), dithiothreitol (DTT; Promega, Madison, WI, USA), and Calcofluor white (10 mg/mL in H2O; fluorescence brightener 28; Millipore Sigma) were prepared and diluted to concentrations indicated in the text.

### Strains and Plasmids Design

All primers and plasmids used in this study are listed in **Supp File 1**. All plasmids generated in this paper underwent whole plasmid sequencing by Plasmidsaurus using Oxford Nanopore Technology with custom analysis and annotation.

*ire1DX* was transformed with the indicated plasmids to produce *ire1DX+IRE1-WT* and other *ire1DX* variants following the protocol described in Sircaik et al., 2021. Briefly, plasmids were cut with NruI to direct insertion into the *C. albicans HIS1* locus. Correct insertion into the locus was confirmed using *IRE1* Comp primers (**Supp File 1**) [36].

CRISPR strains were designed following the protocol in Halder et al., 2019 [92]. Briefly, EuPaGDT (http://grna.ctegd.uga.edu/; Tarleton Research Group, The Center for Tropical and Emerging Global Diseases, The University of Georgia) was used to design small guide RNAs (sgRNA) specific to the gene of interest. The sgRNA sequences and gene-specific homology arms were incorporated into a Gene Drive fragment synthesized by IDT (Integrated DNA Technologies, Inc.; **Supp File 1**). The gene drive was cloned into the *pRS252-Neut5L-CaCas9* backbone using Gibson Assembly, and the resulting plasmid was amplified in *E. coli*. Purified plasmids were sequenced with Plasmidsaurus to confirm correct integration of the gene drive. Plasmids were cut with PacI and transformed into the indicated *C. albicans* strain. Transformation success was assessed using gene drive deletion primers to confirm deletion of the gene of interest (**Supp File 1**).

*pRS313 ScPromoter-CaIRE1* was constructed by amplifying *C. albicans IRE1* (*CaIRE1*) from SC5314 genomic DNA using *CaIRE1* EcoRI primers (**Supp File 1**). The resulting fragment was cut with EcoRI and cloned into a vector containing the *S. cerevisiae IRE1* promoter but no open reading frame (*pRS313-ScIRE1Promoter*; **Supp File 1**) to generate *C. albicans IRE1* under control of the *S. cerevisiae IRE1* promoter. *pRS313-ScIRE1Promoter* was generated by cloning the *S. cerevisiae IRE1* promoter (synthesized by Genewiz, Azenta Life Sciences, South Plainfield, NJ, USA; **Supp File 1**) cut with EcoRI/XhoI into *pRS313*.

### Spot Assays

*C. albicans* cells from overnight cultures were diluted to an OD_600_ of 0.1 and then serially diluted 1:5 in a 96-well plate using appropriate media. Four sequential 1:5 serial dilutions were prepared in the 96-well plate, and a 48-pin replica plater was used to spot the cells onto agar plates containing the indicated concentrations of drugs. Plates were incubated overnight at 30 °C and imaged using a SP Imager (S&P Robotics, North York, ON, Canada).

Quantification either using an accompanying liquid growth assay (**Figure 1**) or using Image J [114] following the protocol described in Petropavlovskiy et al., 2020. Briefly, images were converted to 8-bit format and background noise was subtracted using the Rolling Ball algorithm. A mean grey value was calculated for the background, and grey values of cell spots were measured using the same dilution for each strain. The mean grey value of the background was subtracted from the grey value of each spot, and the resulting values were used for quantification [115]. Specific normalization and statistical analysis information is described in the corresponding figure legends. Quantification was performed using IBM SPSS Statistics version 30.0.0.0 (172) for Windows (IBM Corporation, Armonk, NY, USA).

### Liquid Growth Assays

Overnight cultures of *C. albicans* grown in triplicate were diluted to an OD_600_ of 0.1 and then treated with the indicated drugs. 200 µL aliquots of cells were distributed to a 96-well plate. The plates were placed in an Epoch 2 Microplate Reader (BioTek, Winooski, VT, USA) and OD_600_ readings were measured every 15 minutes for 24 hours. Growth curves were generated and area under the curve was calculated using GraphPad Prism version 10.5.0 for Windows (GraphPad Software, Boston, MA, USA). Statistical analysis was performed using a one-way ANOVA test followed by Tukey’s multiple comparison using IBM SPSS Statistics for Windows, version 30.0.0.0 (172) (IBM Corporation, Armonk, NY, USA)

### Filamentation Assays and Quantification

Cells were grown overnight in triplicate in YPD or SC media. In the morning, cells were diluted to OD_600_ of 0.2 in fresh media and left untreated or treated with 10% fetal bovine serum. Samples were then split into two tubes for growth at both 30°C and 37°C. All samples were incubated for 4 hours. Samples were plated on microscope slides, and imaging was performed using a BioTek Cytation 5 Cell Imaging Multimode Reader.

A protocol for quantifying percent filamentation was developed using a combination of two published protocols [109,116]. **Sample Preparation:** Samples were diluted so that, at a medium magnification, there were no more than ∼100 cells in the field of view. **Image Acquisition:** Using the Cytation5 microscope, three images were taken per replicate for a total of nine photos per treatment condition. Images were taken at random locations throughout the slide, and images with fewer than 10 cells were rejected. The field of view was adjusted as needed to ensure cells and filaments were fully visible and not cut off on the edge. **Categorization of Cells:** For each photo, cells were classified as either round or filamentous. Round cells were defined as being a single round cell, or a round cell with an extension less than twice the diameter of the mother cell body (**Supp Fig. 1A**). Buds are not considered unique cells and are counted as part of the mother cell. Filamentous cells were defined as a mother cell with a filamentous projection at least twice the length of the mother cell body (**Supp Fig. 1B**). **Cell Counting and Statistical Analysis:** Total round and total filamentous cells were counted. For large flocs of cells, the center clump of the floc was excluded from the quantification since the cells become too dense to accurately classify cell type. Percent filamentous cells was calculated for each photo and average percent filamentous cells for each sample was plotted with standard deviation. Statistical analysis was performed using a one-way ANOVA test followed by Tukey’s multiple comparison using SPSS Software.

### RNA Purification and Quality Control

DAY286 and *ire1DX* were cultured overnight in quadruplicate. In the morning, cells were diluted to an OD_600_ of 0.2 and grown for 90 minutes to obtain log phase cells. Cells were then treated with 1.5 μg/mL tunicamycin for two hours at 30°C or with 10% fetal bovine serum for four hours at 37°C. Untreated controls for each strain were grown for two hours at 30°C. Total RNA was isolated using a MasterPure Yeast RNA Purification Kit (LGC Biosearch Technologies). DNase treatment was performed using the DNA-free DNA Removal Kit (Invitrogen). The quality of purified RNA samples was assessed by Bioanalyzer RNA analysis (Agilent; performed by the London Regional Genomics Centre) to ensure an RNA Integrity Number (RIN) of 8 or higher.

### RNA Sequencing

Library preparation and RNA sequencing analysis were conducted by Azenta Life Sciences using an Illumina HiSeq sequencer. The final concentration of total RNA used for library preparation ranged from 37.3-98.5 ng/µL (1.12-2.96 µg). Total RNA was purified using polyA selection and RNA samples were then converted into Illumina TruSeq cDNA libraries. Each sample yielded between 19.4 to 30.1 million 150 bp paired-end sequencing reads. Raw data in FASTQ format and normalized counts were deposited in NCBI’s Gene Expression Omnibus [117] and are accessible through GEO Series accession number GSE308581.

Quality control and trimming was performed using Trim Galore, a wrapper tool around CutAdapt [118] and FastQC (https://www.bioinformatics.babraham.ac.uk/projects/fastqc/). Trim Galore default settings were used to remove Illumina adapter sequences and to remove any reads with a length below 50 bp. Reads were then aligned to the reference genome of *Candida albicans* SC5314 Assembly 22 (NCBI RefSeq assembly GCF_000182965.3; GenBank assembly GCA_000182965.3) using STAR [119]. FeatureCounts was used to count the reads mapped to each gene [120]. Differential expression analysis was performed with DESeq2 [121]. Independent hypothesis weighting was performed with an alpha value of 0.1 and using a comparison that takes into account both strain and treatment [122] (**Supp File 1**). An interaction was added to the model to test if the effect of treatment differs according to strain. K-means clustering based on Euclidean distance was used to cluster significant transcripts in nine and seven groups for the tunicamycin and morphogenesis dataset, respectively (**Supp File 1**). Heat maps were generated using only genes with a p_adj_<0.05 (n=3429 genes for tunicamycin dataset; n=3480 genes for morphogenesis dataset) [123]. Gene ontology analysis for biological processes was performed using ViSEAGO [124]. Gene ontology analysis for cellular compartments (morphogenesis dataset) was performed with GO Enrichment Analysis, powered by PANTHER [125–127] (**Supp File 1**). Data for principal component analysis plots underwent variance stabilizing transformation.

### Hit Validation & GRACE Library

The two clusters containing genes upregulated in an Ire1- and tunicamycin- or serum+37°C-dependent manner (Cluster 2 and 6, respectively) were selected and the top differentially expressed genes from these clusters were determined. The top differentially expressed genes were then compared to strains available in the GRACE Library (Gene Replacement And Conditional Expression). Strains in this library contain mutations that enable a specific gene to be repressed upon treatment with doxycycline [72]. Only genes with mutants available in the GRACE library or its expansion were selected [72,73]. A mini library of 21 strains (tunicamycin) and 20 strains (serum) were constructed (**Supp File 1**). Each contains CaSS1 as a control strain, and each strain was represented in the library in quadruplicate. *HAC1* was added as an additional strain to the tunicamycin library; it was not in the top 20 genes but was of interest given its role in the UPR.

For the tunicamycin mini-library, strains were spotted into liquid YPD and grown overnight at 30°C. Next, the strains were spotted into YPD, YPD + 0.05 μg/mL doxycycline, or YPD + 20 μg/mL doxycycline for induction and grown overnight at 30°C. The lower dose and higher dose of doxycycline induce gene repression for essential and non-essential genes, respectively. The following day, strains from each condition were spotted into YPD media containing the same doxycycline treatment with or without 1.5 μg/mL tunicamycin and grown overnight at 30°C. Plates were then mixed briefly prior to measuring the endpoint OD_600_. Data was analyzed as described by Lee et al., 2022. Median absolute deviation (MAD) was calculated for the -tunicamycin and +tunicamycin conditions and plotted. Genes in the lower left and right quadrants were identified as hits [74]. Several genes falling slightly outside of these quadrants were still considered as genes of interest.

For the top 20 genes from morphogenesis Cluster 6, the PathoYeastrack Rank Genes by Transcription Factor function was used to search for transcription factors that regulate the group of 20 genes [90]. The search criteria for this included: Documented regulations were filtered by DNA binding or expression evidence, and all expression evidence was included (TF acting as activator, TF acting as inhibitor, and regulations without association information). All transcription factors were checked for, and no other criteria were specified. The full table of results is reported in **Supp File 1**. Transcription factors were categorized as regulating morphogenesis, biofilm formation, adhesion, or another function based on GO annotations and mutant phenotype data available in the Candida Genome Database [91].

### Potential RIDD Target Determination

This protocol was based on analysis performed in Kimmig et al., 2012, for *S. pombe [58]*. For all genes detected in our RNA sequencing (n>6,000 genes), the effect of strain in the treated samples (tunicamycin or serum+37°C), and the treatment effect in the wild-type strain were calculated. Genes with fold changes with p_adj_<0.05 for both variables were plotted and considered for further analysis (**Fig. 5A/B**; n=2685 genes tunicamycin, n=1773 genes morphogenesis). For the tunicamycin treatment, 30 genes with a decrease in expression greater than 1.5-fold associated with both strain and tunicamycin treatment were considered (**Supp File 1**). For the serum+37°C treatment, 23 genes with a decrease in expression greater than 1.0-fold associated with both strain and serum+37°C were considered (**Supp File 1**). The Candida Genome Database was used to determine protein product, cell component, and presence of a transmembrane domain for each gene [91]. The CGD definition of cell surface is any protein localized to the external part of the cell wall and/or plasma membrane.

### qRT-PCR

Overnight cell cultures were grown in quadruplicate. In the morning, cultures were diluted 1:10 in appropriate media and grown for 90 minutes to log phase, then treated with 1.5 μg/mL tunicamycin for two additional hours at 30°C or with 10% fetal bovine serum for four additional hours at 37°C. Untreated cells were grown an additional two hours at 37°C to serve as a control. RNA purification and DNase treatment were performed as described above. cDNA libraries were prepared using 2.5 μg total RNA using the SuperScript IV VILO Master Mix (Thermo Fisher Scientific) according to the manufacturer’s instructions. Primers used for qRT-PCR can be found in **Supp File 1**. *ACT1* was used as a housekeeping gene for all *C. albicans* genes, and qRT-PCR was performed using a QuantStudio 3 Real-Time PCR system (Thermo Fisher Scientific) using the comparative Ct method of analysis [128]. Unless otherwise stated, all ΔCt values were normalized to the ΔCt of the untreated control strain to obtain ΔΔCt values.

### *HAC1* Splicing

Overnight DAY286 cultures were diluted to OD_600_ 0.2 and grown to log phase. Tunicamycin and fetal bovine serum treatments were administered as described in the RNA Purification methods section. For the splicing time course with serum, cells were immediately removed after addition of serum to serve as the 0 minute time point. Additional aliquots of cells were removed at the indicated time points post-treatment, after which they were washed with cold water and frozen until RNA purification.

RNA purification and DNase treatment were performed as described above. RT-PCR was performed using *HAC1* splicing primers (**Supp File 1**) with the SuperScript IV One-Step RT-PCR System (Thermo Fisher Scientific). Samples were run on 4% agarose gels and imaged using a Gel Doc XR+ Imager (Bio-Rad). Gel quantification was performed following the methods described in Petropavlovskiy et al., 2020 to obtain mean grey values (MGV) for each band. The MGV ratio of spliced *HAC1* to total *HAC1* was determined and one-way ANOVA with Tukey’s multiple comparison was performed using SPSS.

## Acknowledgements

PL is supported by an NSERC Discovery Grant (RGPIN-2022-05267), CIHR Project Grants (PJT 168882 and ARB 192062). PL is also supported by a Western CIHR Accelerator grant as well as a Western CIHR Reapplication Program grant. VD is supported by an NSERC Discovery Grant (RGPIN-2024-04932). RSS holds the Canada Research Chair in Microbial Functional Genomics and Synthetic Biology. SSC holds an Ontario Graduate Scholarship. BL was the recipient of an NSERC Undergraduate Summer Research Award. The *ire1DX* strain and the *IRE1-WT*, *IRE1-ND*, and *IRE1-KD* plasmids were obtained from Sneh Panwar (Jawaharlal Nehru University). The original *pDDB78* vector was a gift from Aaron Mitchell (University of Georgia). We thank MERK and Leah Cowen (University of Toronto) for making the GRACE library available.

