## Supplemental Figures for "Distinct Ire1-driven transcriptional responses control morphogenesis in *Candida albicans*"

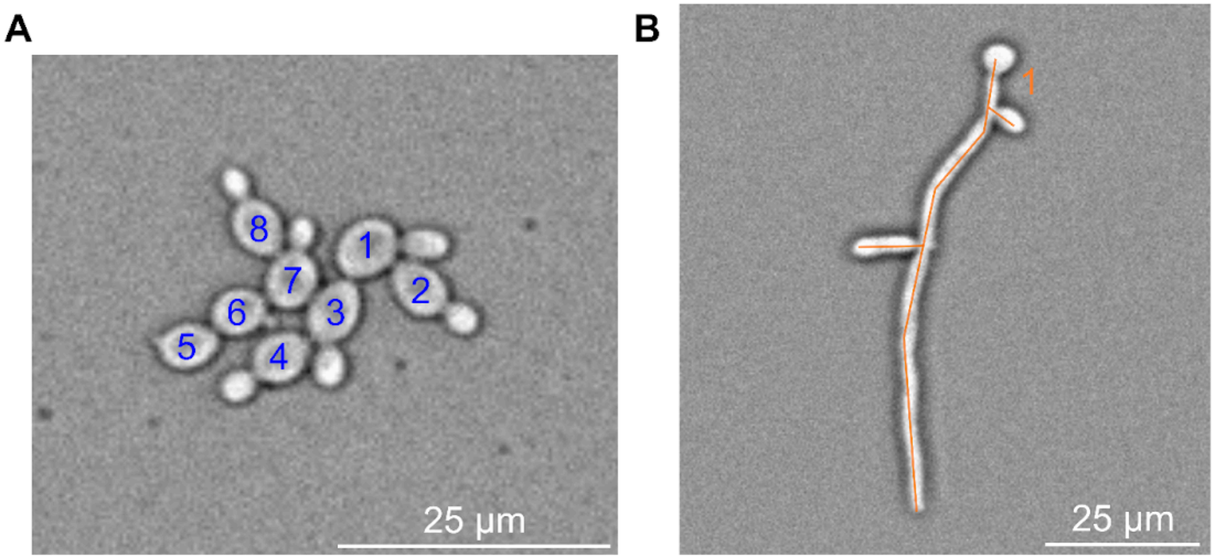


**Supplemental Figure 1. Filament quantification method.** Cells were classified as round or filamentous for quantification of percent of cells filamenting. **A.** Round cells, numbered in blue. Some round cells have buds, which are not considered unique cells during counting. **B.** Filamentous cell, outlined in orange.

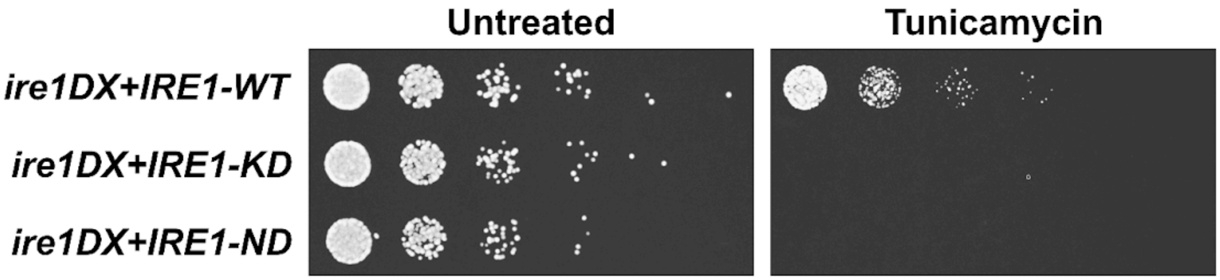


**Supplemental Figure 2. Ire1 kinase and nuclease domains are required for growth in the presence of tunicamycin.** *C. albicans ire1DX* cells complemented with wild-type (WT) *IRE1*, kinase-dead (KD) *IRE1*, and nuclease-dead (ND) *IRE1* were grown on agarose plates with 1.5 µg/mL tunicamycin. Images acquired after 24 hours.

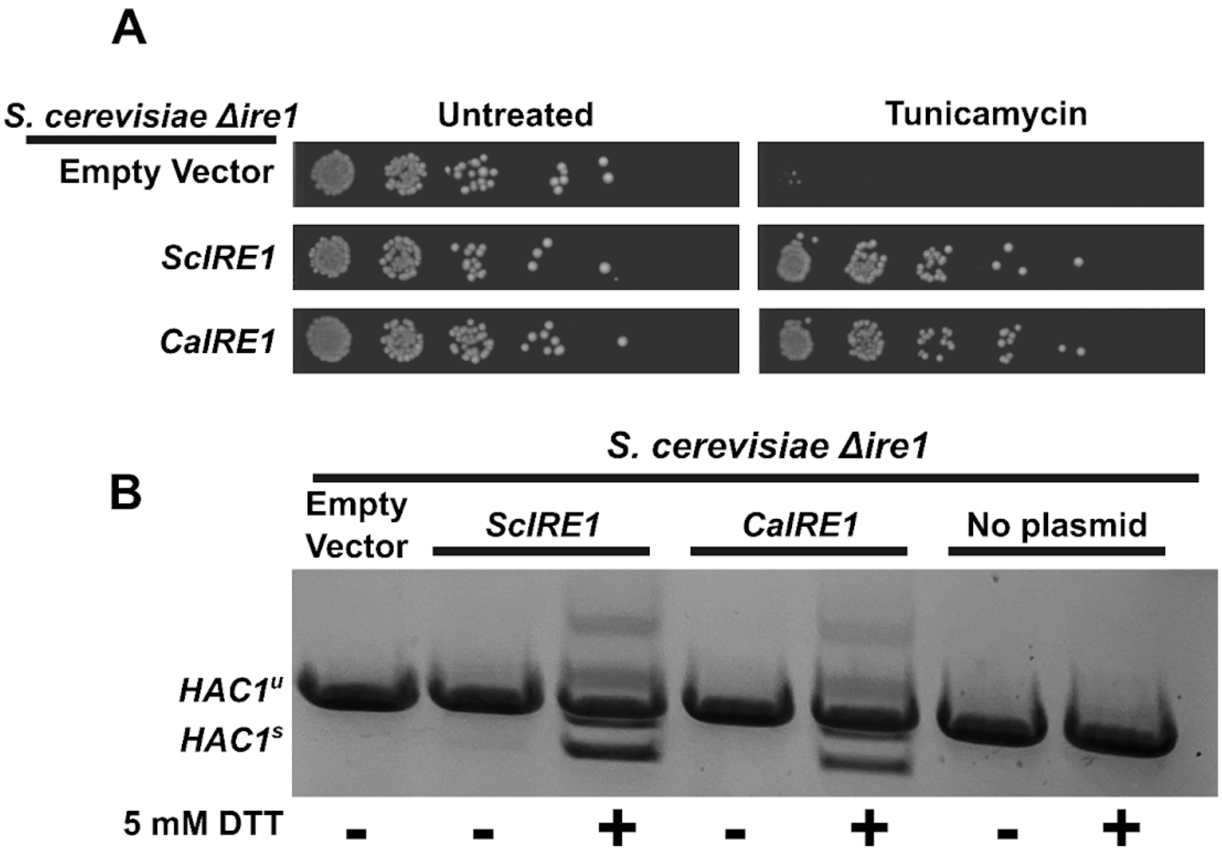


**Supplemental Figure 3. *C. albicans IRE1* complements the *IRE1* function of *S. cerevisiae*.** **A.** *S. cerevisiae ire1Δ* cells containing the indicated plasmids were grown on agar plates with 1.5 μg/mL tunicamycin. The empty vector is *pRS313*, *ScIRE1* is *pRS313-ScIRE1*, and *CaIRE1* is *pRS313-ScPromoter-CaIRE1*. Images acquired after 48 hours. **B.** The same strains were treated with 5 mM DTT for 40 minutes and total RNA was purified. RT-PCR was performed using primers flanking the *HAC1* intron and DNA was run on a 1% agarose gel.

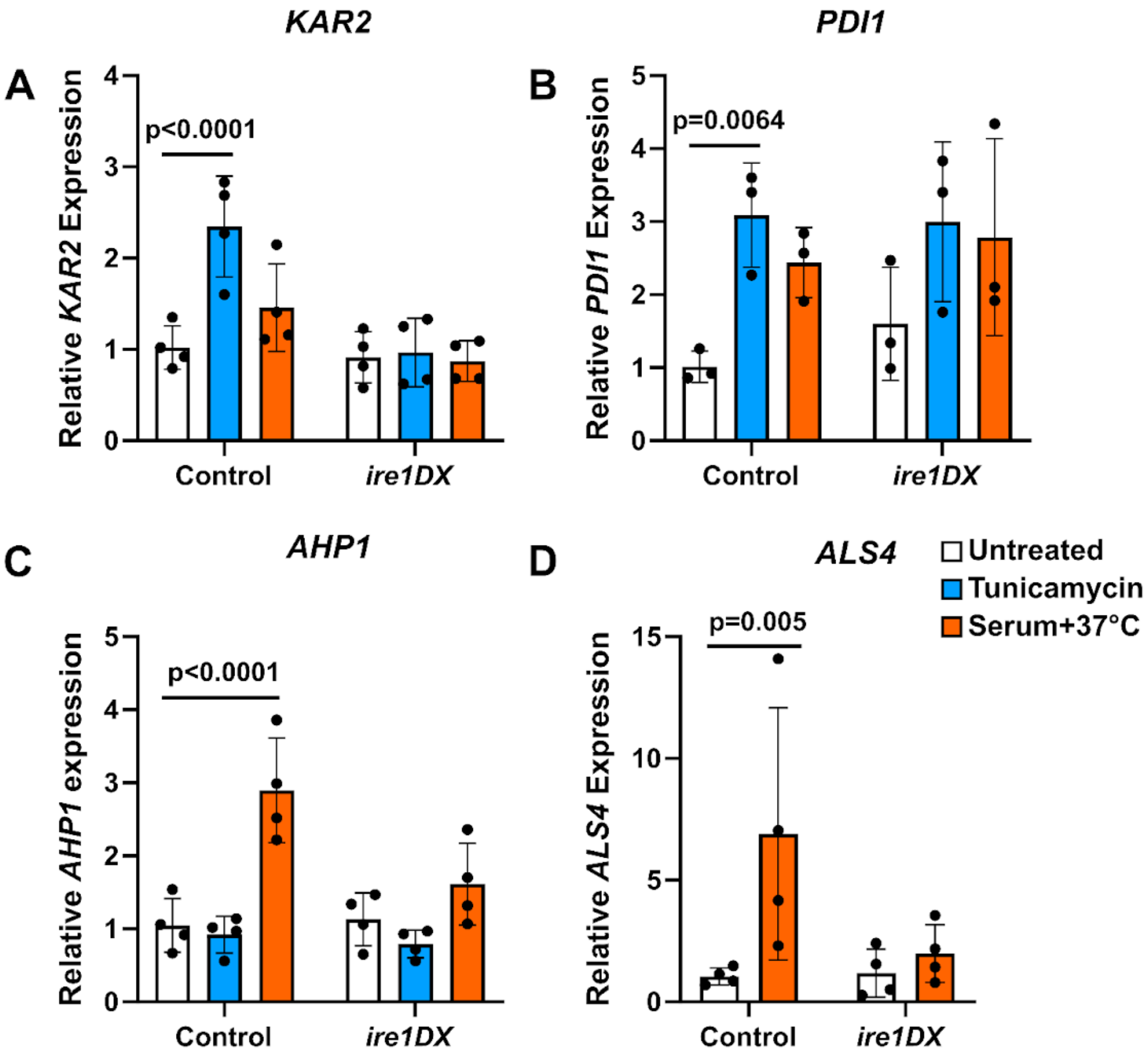


**Supplemental Figure 4. RT-qPCR validation of RNA sequencing hits.** Relative gene expression for **A.** *KAR2*, **B.** *PDI1*, **C.** *AHP1*, and **D.** *ALS4* in control (DAY286) and *ire1DX* strains. Mean±SD shown. One-way ANOVA with Tukey's multiple comparisons performed in GraphPad Prism. n=3 for *PDI1* and n=4 for *KAR2, AHP1*, and *ALS4*. Legend applies to all graphs.

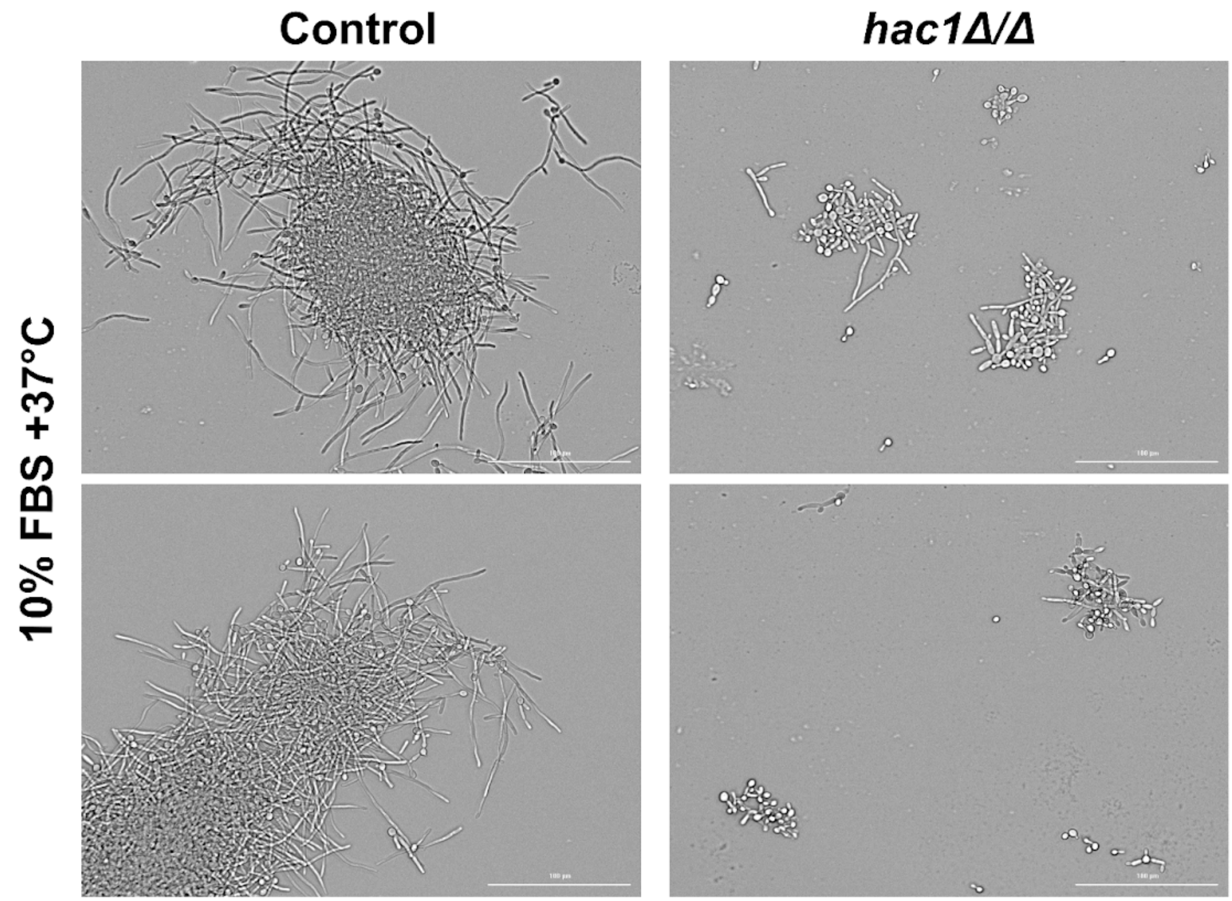


**Supplemental Figure 5. *C. albicans* cells lacking *HAC1* do not form large flocs.** Control (DAY286-Cas9) and *hac1Δ/Δ* strains were grown in filamentation-inducing 10% fetal bovine serum (FBS) for four hours at 37°C. Images obtained using the Cytation5 cell imaging multi-mode reader. Scale bar = 100 µm.

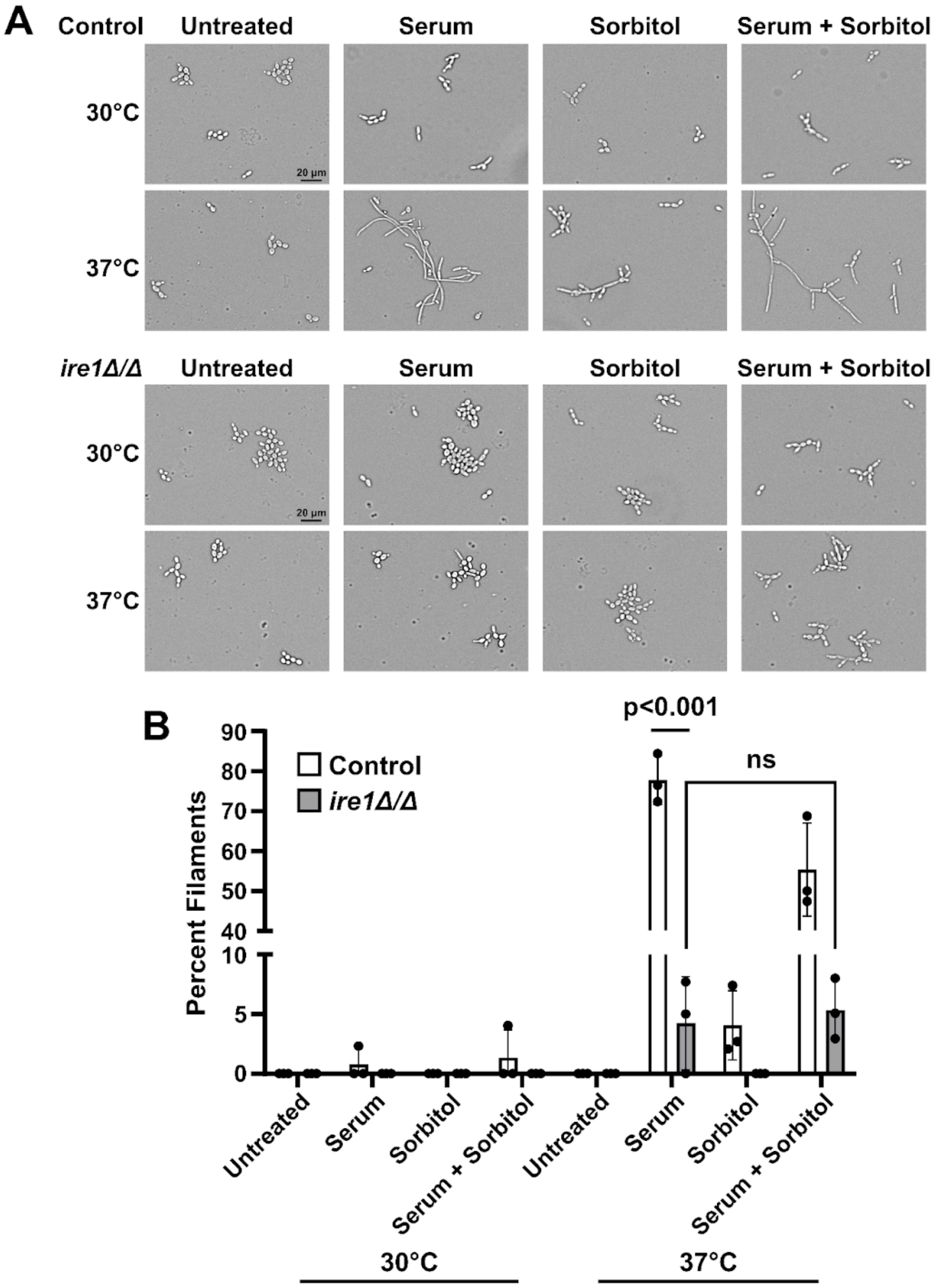
A
Control
Untreated
Serum
Sorbitol
Serum + Sorbitol
30°C
37°C
20 µm
ire1Δ/Δ
Untreated
Serum
Sorbitol
Serum + Sorbitol
30°C
37°C
20 µm
B
p<0.001
ns
Control
ire1Δ/Δ
Percent Filaments
90
80
70
60
50
40
10
5
0
Untreated
Serum
Sorbitol
Serum + Sorbitol
Untreated
Serum
Sorbitol
Serum + Sorbitol
30°C
37°C

**Supplemental Figure 6. Sorbitol does not rescue morphogenesis in *IRE1* deficient cells.** **A.** DAY286-Cas9 (control) and *ire1Δ/Δ* cells were grown in filamentation-inducing 10% fetal bovine serum and 1 M sorbitol for four hours at 37°C and at 30°C as a control. Images obtained using the Cytation5 cell imaging multi-mode reader. Scale bar = 20 μm, applies to all images. **B.** Filamentation was quantified using photos acquired from randomly selected portions of each slide, and round and filamentous cells were counted by hand for each photo. Percent filaments was calculated and mean±SD was plotted. n=3.
